# Nucleic acid amplification by a transparent graphene Visual-PCR chip and a disposable thermocycler

**DOI:** 10.1101/724245

**Authors:** Guozhi Zhu, Miao Qiao

## Abstract

Polymerase chain reaction (PCR) is a method widely used to amplify trace amount of nucleic acids. It needs a process of thermocycling (repeated alternation of temperature). Traditional thermocycler relies on bulk size of metal block to achieve thermocycling, which results in high cost and the lack of portability. Here, a PCR chip made of graphene Transparent Conductive Films (TCFs) was employed. The thermocycling of the chip was fulfilled by a temperature programed microcontroller and a cooling fan under a low driving voltage (12V). A 35 cycles PCR was accomplished within 13 minutes using the chip and the thermocycler. The transparency of the graphene PCR chip enables the PCR reaction to be visually monitored by naked eye for a color change. The PCR chip and the thermocycler have a low cost at $2.5 and $6 respectively, and thus are feasible for Point-of-care testing (POCT) of nucleic acids in a disposable manner. The whole platform makes it possible to perform a low-cost testing of nucleic acids for varieties of purposes outside of laboratories or at resource limited locations.

## INTRODUCTION

Polymerase chain reaction (PCR) (Saiki, et al., 1988) as the basic method of Molecular Diagnostics (MDx) has been widely used not only in academic research but also in applications of clinic, food safety, veterinary, etc (Ishmael, et al., 2008). PCR relies on repeated cycles of heating and cooling (thermocycling) to amplify trace amount of nucleic acids. Modern thermocycler commonly uses Peltier elements to fulfill thermocycling. Quality Peltier element is made of costly metal, such as silver. The amplified PCR products are frequently determined by fluorescent analysis, which makes thermocycler to be even more costly (Navarro, et al. 2015).

Graphene is two-dimensional, hexagonal lattice of carbon atoms (Geim, 2007; Geim, 2009). Graphene exhibits outstanding electrical conductivity, transparency and flexibility, which make graphene transparent conductive films (TCFs) to be an excellent material for touch screen and varieties of applications (Yu, et al, 2017; Zhang, et al., 2019). Graphene TCFs show very fast heating temperature response even under a low driving voltage (Kang, et al., 2011; Sui, et al., 2011). In addition, the low heat capacity of graphene makes its film to be cooled down rapidly (Kim, et al., 2018; Zhang, et al, 2017). These years witness the large scaled production capacity of graphene TCFs, which results in its dramatic price drop.

“Hyper-local is the future of healthcare” (Bertolini, et al., 2017). Point-of-care testing (POCT), such as handheld blood glucose meter, dramatically provides convenience and represents the trend of healthcare. In this respect, POCT MDx is still at its earlier stage and an ideal POCT MDx platform with low cost and portability is missing. The current report presents a low-cost and portable POCT MDx platform.

Taking advantage of the low cost and fast heating/cooling of graphene TCFs, a PCR chip was made here. The thermocycling of the graphene PCR chip was successfully achieved to drive a PCR reaction. Fast and sensitive PCR reactions were accomplished. Integrated with the non-fluorescent colorimetric PCR technology (Zhu, et al. 2018), the platform successfully demonstrated the visual detection of PCR reaction in the transparent graphene PCR chip. The advantages of low cost, portability, fast speed and sensitivity enable the platform to be suitable for diagnostic applications in POCT or resource-limiting areas.

## MATERIAL AND METHODS

### Graphene TCF, PCR reaction well and PCR kit

Commercial graphene TCFs were purchased from Changzhou Erwei Tansu Technology Co. Two graphene TCFs were used (catalog number JR029-H and JR029-E, $6 for each). Both of the graphene TCFs have 120 × 141 mm in size, 0.25mm in thickness. Both are transparent and have 83% efficiency from electricity to heat. JR029-H has a 2.8Ω electrical resistance, whereas JR029-E measures a 5.8Ω electrical resistance.

Polypropylene film wells were employed as PCR reaction wells in graphene PCR chip. Polypropylene film is thin enough to ensure fast heat transfer. Either home-made wells or wells of NGPCR 96-well microplate (MBS, Netherlands) were used. The home-made wells were prepared by heat-pressing small stainless metal balls against a sheet of polypropylene film (0.05mm).

A commercial Visual-PCR kit (catalog number GM7099, GM Biosciences) for testing GFP tag was employed to prepare PCR reaction solution following the kit instruction. The PCR reaction solution contains a colorimetric dye to monitoring PCR reaction. The dye is visually detectable colored dye which changes its original violet-purple color to blue upon PCR amplification. Naked eye can easily discriminate the color change.

### Setup of graphene PCR chip

Transparent graphene PCR chip comprises of graphene TCFs, PCR reaction wells and temperature sensor. Two graphene TCFs sandwich PCR reaction wells and a temperature sensor to give a complete graphene PCR chip reported here. An exemplary PCR chip was prepared as followed. A small piece (120 × 34mm) of graphene TCFs was cut from either JR029-E or JR029-H graphene TCFs, which gave a measured 35Ω and 17Ω electrical resistance. Folded the piece of film in half, attached its two ends together, and stapled its edges to give size of 58 × 34mm. 5ul PCR reaction solution were loaded into polypropylene film well. The well was then heat sealed to the other polypropylene film on top to completely insulate the PCR reaction solution. The sealed wells were then placed between two graphene TCFs. A K-type thermocouple was inserted into the two graphene TCFs at a location close to the PCR reaction wells (within 1cm distance) to monitor the chip temperature. Thus, the exemplary PCR chip has two graphene TCFs which sandwich PCR reaction wells and a temperature sensor, as illustrated in Fig 1A. Photos of the transparent graphene PCR chips were shown in Fig 2 and Fig 3.

**Fig 1.**
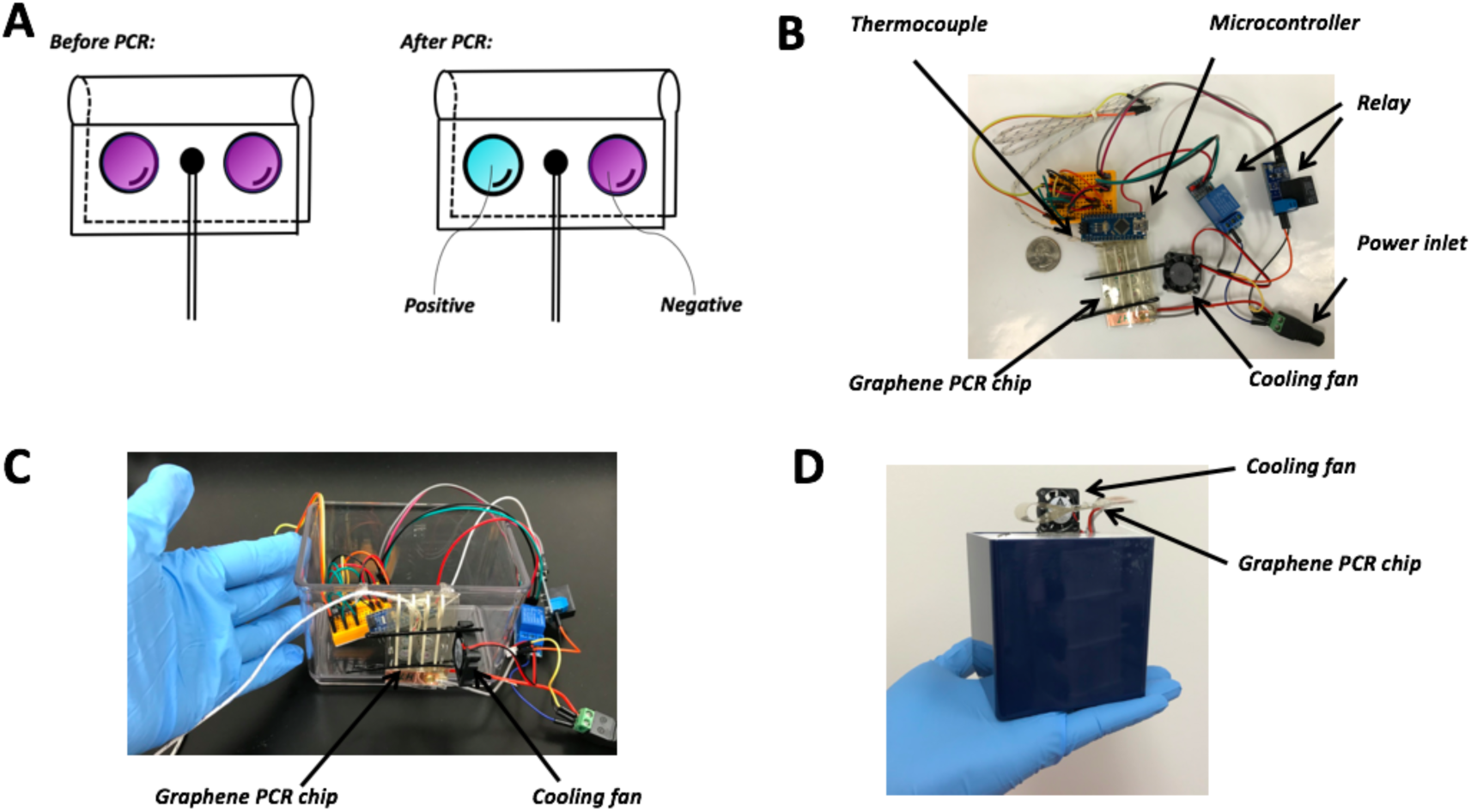
Illustration of the graphene Visual-PCR chip and setup of the low-cost disposable thermocycler. (A) Illustration of a transparent graphene Visual-PCR chip before and after PCR. The graphene PCR chip is made by folding a graphene TCF that sandwich one thermocouple and two reaction wells. Before PCR, the PCR solution in reaction wells are purple color. After PCR, the reaction wells give different colors contingent upon the presence or absence of templates. Positive reaction changes to sky-blue, whereas negative reaction keeps its original purple color. The color change can be observed directly by naked-eyes in a real-time manner. (B) Major components of the thermocycler include a transparent graphene PCR chip with a thermocouple inserted inside, a microcontroller (Arduino Nano electronic controller), a cooling fan, two relays controlling the power to the PCR chip and cooling fan respectively. Temperature programed microcontroller automatically controls time/duration to power on/off the PCR chip and cooling fan. (C) and (D) Pictures of the thermocycler prototype in an open and closed format. Both the graphene PCR chip and cooling fan were attached to the sidewall of a box. The graphene PCR chip is positioned vertically to the cooling fan.

**Fig 2.**
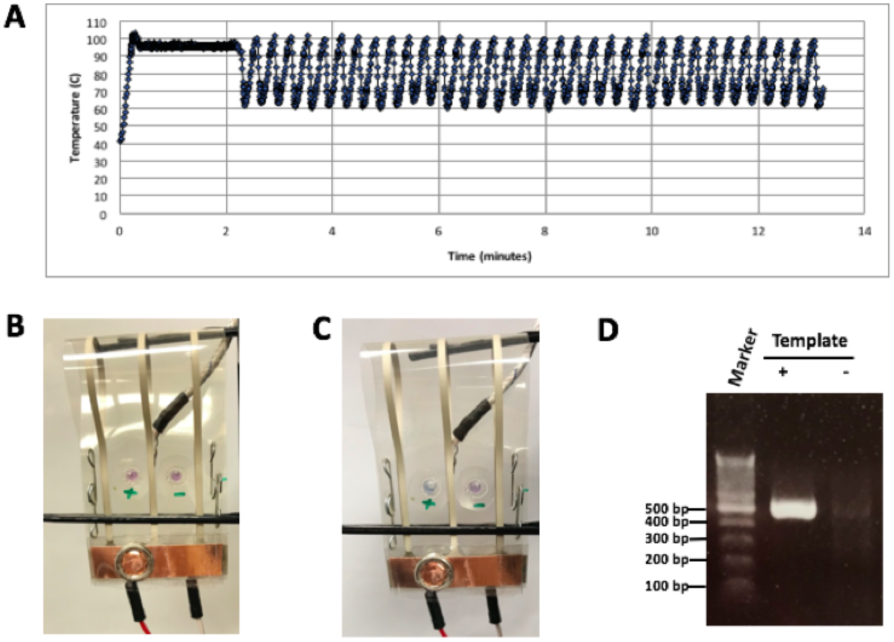
Fast PCR performed in graphene Visual-PCR chip. (A) Temperature plot of a complete PCR reaction (35 cycles in approximately 13min). (B) The photograph of the graphene Visual-PCR chip prior to PCR. A piece of graphene TCFs JR029-E was used to make the chip. (C) The photograph of the same PCR chip after PCR reaction. (D) The gel electrophoresis of the PCR reactions of C.

**Fig 3.**
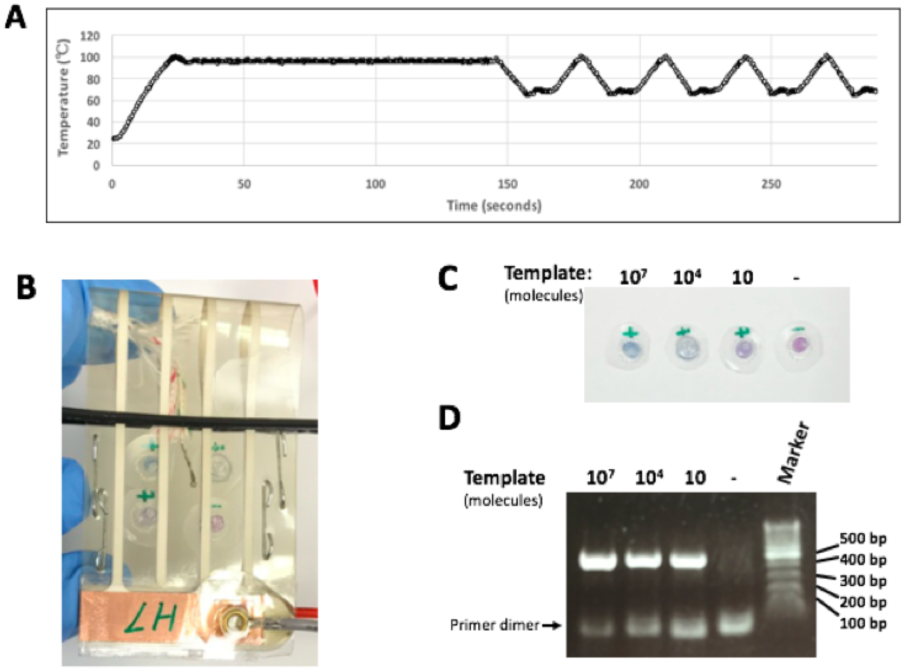
Sensitive PCR performed in graphene Visual-PCR chip. (A) The expanded view of the temperature plot that covers the initial denaturation step and the first 5 cycles. (B) Photograph of the graphene PCR chip after PCR completion (50 cycles). (C) The 4 reaction wells were taken out of the PCR chip and photograph was taken. They showed different colors. (D). The PCR products of the 4 reactions were subjected to agarose gel electrophoresis.

### Setup of thermocycler and PCR condition

The thermocycler is made of microcontroller, cooling fan, relays, etc., as illustrated in Fig 1B. The microcontroller includes an Arduino Nano board and a thermocouple breakout amplifier (AD595) connected with a K-type thermocouple from graphene PCR chip. The thermocouple breakout amplifier reads the temperature from the thermocouple and transfers the data to Arduino Nano board. The Arduino Nano reads the data from the amplifier and output it to a computer.

Both the graphene PCR chip and the cooling fan were connected to the microcontroller through a relay respectively. The cooling fan was positioned at the one flank of the PCR chip so that the surface of both sides (front and rear) of PCR chip obtains sufficient air circulation.

Both the PCR chip and the cooling fan are automatically controlled by the microcontroller. When heating the PCR chip, the graphene TCFs are powered on and the cooling fan is powered off. When cooling the PCR chip, the graphene TCFs are powered off and the cooling fan is powered on. Based on the chip temperature sensed in a real-time manner, the microcontroller is programed in such a way that it automatically determines the time/duration to turn on/off the power to graphene TCFs, and the time/duration to turn on/off the cooling fan. The microcontroller was loaded with PCR program: 94°C for 2 minutes followed by 35-50 cycles of 94°C for 4 seconds and 70°C for 7 or 10 seconds.

### Gel electrophoresis analysis

After PCR amplification, the PCR reaction was subjected to agarose gel electrophoresis. 1.5% pre-cast gel (Lonza, Rockland, ME) was run at 110V for 10-20 minutes. The resulting gel images were captured by a camera.

## RESULTS

Transparent graphene TCFs were used here to replace traditional metal Peltier element to fulfill the thermocycling. A transparent graphene PCR chip consists of transparent graphene TCFs which sandwich PCR reaction wells and a temperature sensor. Visual-PCR technology was employed to take advantage of the transparency of the chip. The color of PCR reactions in the chip was visually identified by naked eyes. The resulted graphene Visual-PCR chip is illustrated in Fig 1A.

A simple thermocycler was constructed, whose components are shown in Fig 1B. As shown, the thermocycler mainly contains a microcontroller, a cooling fan, two relays and a thermocouple. A temperature programed microcontroller senses the chip temperature through the thermocouple, and then automatically controls the power to a graphene PCR chip for heating and the power to a cooling fan for cooling. The prototypes of the thermocycler in an open and closed format are shown as Fig 1C and 1D.

The performance of the whole system was then tested. The microcontroller was loaded with PCR program: 94°C for 2min followed by 35 cycles of 94°C for 4 seconds and 70°C for 7 seconds. A graphene Visual-PCR chip was constructed as shown in Fig 2B. The graphene TCFs were cut from JR029-E and powered with 19.5V voltage through a relay. A CPU cooling fan (DC12V, 0.51A, 8cm × 8cm × 2.5cm in size) was powered by a 12V voltage for cooling the graphene Visual-PCR chip. 5ul Visual-PCR reaction solution with/without 10^8^ copies of templates (GFP gene containing plasmid) were included in reaction wells. The thermal profile of the graphene Visual-PCR chip while running the PCR program was shown in Fig 2A. It was observed that the reaction well with templates changed its color at 25 cycles. After PCR reaction of 35 cycles, the chip was pictured and shown in Fig 2C. The PCR reactions were then subjected to gel electrophoresis shown in Fig 2D.

As shown, the temperature at the initial denature step was maintained at 94°C steadily (Fig 2A). 35 cycles only took approximately 13 minutes. Each cycle took an average of 19.0 seconds. Of the 19.0 seconds, heating the chip from 70°C to 95°C took only an average of 5.2 seconds, and cooling from 95°C to 70°C took only an average of 2.7 seconds.

Before PCR, the PCR solution showed purple color (Fig 2B). After PCR, the reaction with template (left) changed to sky-blue color and the non-template one (right) kept its original purple color (Fig 2C). Consistently, gel electrophoresis (Fig 2D) showed a desired 500bp amplification band for positive reaction but not for negative reaction. It verified that the reaction with blue color had specific amplification, whereas the reaction with purple color did not have specific amplification. As shown in Fig 2, fast PCR was accomplished by the graphene Visual-PCR chip.

The sensitivity of the system was tested thereafter. The microcontroller was loaded with PCR program: 94°C for 2 minutes followed by 50 cycles of 94°C for 4 seconds and 70°C for 10 seconds. A piece of graphene TCF JR029-H was used to make a new graphene Visual-PCR chip. A mini-cooling fan (DC12V, 0.07A, 2.5cm X 2.5cm x 0.7cm in size) was employed. Both the PCR chip and the mini-cooling fan were powered by a same 12V power. Four reactions with different copies of templates were included in the graphene Visual-PCR chip. The temperature profile is shown at Fig 3A, and the photo of the chip after PCR is shown as Fig 3B. The four reactions were taken out of the chip and pictured as Fig 3C, and then subjected to electrophoresis as Fig 3D.

Fig 3A shows the annealing/extension step can be maintained at 70°C to some extend instead of a sharp temperature diving in Fig 2A. This is due to a smaller cooling fan utilized in this experiment.

Fig 3B and 3C clearly shows that all of the reactions with templates of 10^7^, 10^4^ and 10^1^ copies have a visible color change. The result of gel electrophoresis shown in Fig 3D re-confirmed the specific amplifications for all of them. According to the experiment of Fig 3, the sensitivity of the whole system can reach to 10 copies. Sensitivity lower than 10 copies has not been tested yet.

The thermocycler used here costs about $21. Its components can be obtained at lower prices shown in Table 1. A similar thermocycler only costs as low as $6. Briefly, an Arduino Nano microcontroller can be obtained for about $2 from AliExpress. A K-type thermocouple amplifier MAX31855 with similar function to AD595 can be obtained for $1.28 from AliExpress. Thermocouple, mini-cooling fan and relays are common electronics with a cost in a range of $0.5- $2 for each. The low cost of thermocycler makes it feasible to be used in a disposable manner.

**Table 1.**
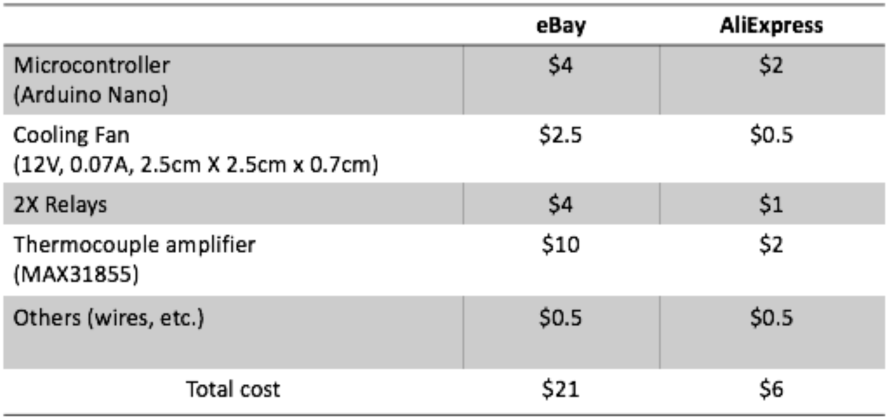
Cost breakdown for a thermocycler. The prices from eBay and AliExpress are shown here.

The graphene PCR chip costs $2.5 ($1.5 for graphene TCFs and $1 for K-type thermocouple). Therefore, the total cost of the PCR chip and thermocycler can be lower than $9.

## DISCUSSION

The current report presents a handheld POCT MDx platform with low-cost, low driving voltage (12V) and 8W power consumption. It makes it possible for people to perform PCR testing at different locations for varieties of purposes.

Several improvements were presented here to make such a PCR platform. Firstly, low cost graphene TCFs were used to replace traditional expensive heavy metal Peltier element, and the graphene TCFs were integrated into a PCR chip. Secondly, Visual-PCR technology was employed so that PCR signals can be visually detected. In third, a very simple and low-cost thermocycler was constructed.

Fast heating and fast cooling are superb characteristics of graphene TCFs. The system takes full advantages of these merits. Both fast thermocycling of the graphene PCR chip and fast PCR reaction were successfully accomplished, as shown in Fig 2.

Visual-PCR provides an easy way to detect signals. The color development during PCR reaction is passed through the transparent graphene PCR chip, which enables real-time color detection and quantification of target nucleic acids. Comparing to traditional fluorescent analysis, colorimetric analysis does not need expensive optoelectronics and filters. Therefore, utilization of Visual-PCR makes the thermocycler as simple as possible. The color monitoring can be further fulfilled by a smartphone App in the future. The App can not only provide an objective reading but also quantify target nucleic acids by real-time data acquisition.

Heating speed of the graphene PCR chip is dependence on the surface resistance of graphene TCFs and the power voltage to drive it. 19.5V voltage has faster heating speed than 12V voltage. Cooling speed can be fine-tuned by choosing suitable cooling fan. For example, a large cooling fan (12v, 0.6A) resulted in faster cooling, whereas a mini-cooling fan (12v, 0.07A) resulted in a slower cooling speed and a stable temperature in extension step. If time is the concern in some scenarios, higher voltage and powerful cooling fan are recommended to speed up the PCR. The performance of the platform can be tailored accordingly to make it more versatile.

The thermocycler here has a low-cost due to its simple structure. A microcontroller, a cooling fan and two relays constitute the majority of thermocycler. However, there are more rooms to reduce the cost further. For example, the relays can be further replaced by bipolar transistors reducing the price to half. A custom microcontroller integrating multiple elements into a single board can further reduces its cost. Therefore, it is very likely to produce a thermocycler for cost of $4-$5 in total.

It is noted that graphene TCF integrating PCR chip was reported previously in a format of convective flow PCR by Chung’s group (Chung, et al, 2014). In their design, graphene TCF kept at stable temperature, and convective flow of PCR solution between two temperature zones was induced to drive PCR reaction. The sensitivity of the graphene convective PCR chip is unknown. In contrast, graphene TCFs in this report are thermocycling instead of keeping at stable temperature. The outstanding features of fast heating and cooling of graphene TCFs are fully utilized here. Sensitive PCR reaction was accomplished and 10 copies were successfully detected (Fig 3).

Traditionally, thermocycler is expensive and it is sold separately from PCR kits. However, the thermocycler reported here can be disposable due to its low cost. In the future, an all-in-one PCR kit includes both the disposable thermocycler and PCR chips. PCR program optimized for a typical pathogen is pre-loaded into the disposable thermocycler. And such an all-in-one kit provides multiple tests for a specific pathogen with an affordable price.

Based on the platform presented here, many improvements on hardware can be done in the future, such as, to reduce thermocycler size to business card size, to integrate a digital screen and provide a friendly user interface, to integrate a WiFi chip, etc. The digital screen functions to report the progress of PCR program and any operational errors such as incorrect connection of chip, unexpected heating/cooling speed, etc. WiFi chip is to wirelessly communicate the thermocycler with smartphone, so that smartphone can initiate/stop the reaction, monitor the color change, upload data to cloud server for a detail analysis and draw a history graphs, etc.

A transparent graphene Visual-PCR chip and a disposable thermocycler are developed here. The whole system provides a platform for POCT MDx with a 12V power input, which make it possible for people to perform PCR testing at a convenient location on pathogens, microorganism, virus, allergens, transgene, genotyping, etc. The platform is low cost without sacrificing its sensitivity and speed. It will have impact on many applications such as clinic, food safety, water/drinking pathogens, veterinary, environmental allergen testing, etc.

